# NMD targets experience deadenylation during their maturation and endonucleolytic cleavage during their decay

**DOI:** 10.1101/2023.09.29.560204

**Authors:** Marcus J. Viscardi, Joshua A. Arribere

## Abstract

Premature stop codon-containing mRNAs can produce truncated and dominantly acting proteins that harm cells. Eukaryotic cells protect themselves by degrading such mRNAs via the Nonsense-Mediated mRNA Decay (NMD) pathway. The precise reactions by which cells attack NMD target mRNAs remain obscure, precluding a mechanistic understanding of NMD and hampering therapeutic efforts to control NMD. A key step in NMD is the decay of the mRNA, which is proposed to occur via several competing models including deadenylation, exonucleolytic decay, and/or endonucleolytic decay. We set out to clarify the relative contributions of these decay mechanisms to NMD, and to identify the role of key factors. Here, we modify and deploy single-molecule nanopore mRNA sequencing to capture full-length NMD targets and their degradation intermediates, and we obtain single-molecule measures of splicing isoform, cleavage state, and poly(A) tail length. We observe robust endonucleolytic cleavage of NMD targets *in vivo* that depends on the nuclease SMG-6 and we use the occurence of cleavages to identify several known NMD targets. We show that NMD target mRNAs experience deadenylation, but similar to the extent that normal mRNAs experience as they enter the translational pool. Furthermore, we show that a factor (SMG-5) that historically was ascribed a function in deadenylation, is in fact required for SMG-6-mediated cleavage. Our results support a model in which NMD factors act in concert to degrade NMD targets in animals via an endonucleolytic cleavage near the stop codon, and suggest that deadenylation is a normal part of mRNA (and NMD target) maturation rather than a facet unique to NMD. Our work clarifies the route by which NMD target mRNAs are attacked in an animal.

## INTRODUCTION

Nonsense-mediated mRNA decay (NMD) is a post-transcriptional surveillance system that targets and degrades mRNAs containing premature termination codons (PTCs), which would otherwise produce truncated, non-functional, and/or toxic proteins (reviewed in (Kurosaki et al. 2019)). NMD plays a crucial role in human health by degrading mRNAs containing PTCs, which are found to drive diverse diseases (Mort et al. 2008). NMD also regulates the abundance of ∼5-20% of endogenous transcripts in animals, making NMD broadly relevant for understanding gene expression (Ramani et al. 2009; Muir et al. 2018; Rehwinkel et al. 2005; Mendell et al. 2004; Wittmann et al. 2006; Kim et al. 2022).

While it is known that NMD degrades targeted mRNAs, multiple competing models exist to describe how the mRNAs are attacked in animals, which feature endonucleolytic cleavage, deadenylation, and/or decapping. In the endonuclease model, the metal-dependent endonuclease domain of SMG-6 first cleaves NMD targets (Gatfield et al. 2003; Glavan et al. 2006; Eberle et al. 2009). The essential role of SMG-6 in NMD function is supported by several studies (Baird et al. 2018; Alexandrov et al. 2017; Zhu et al. 2020; Zinshteyn et al. 2021; Kim and Modena et al. 2022; Colombo et al. 2017; Boehm et al. 2021), and challenged by others (Loh et al. 2013; Jonas et al. 2013; Metze et al. 2013; Huth et al. 2022). In an alternative to SMG-6-mediated cleavage, deadenylation of NMD targets occurs via SMG-5 and SMG-7 (Loh et al. 2013; Chen and Shyu 2003; Lejeune et al. 2003; Yamashita et al. 2005). SMG-7 associates with the deadenylase complex and elicits mRNA decay in tethering assays (Loh et al. 2013), and NMD-targeted mRNAs experience deadenylation in mammalian cells (Chen and Shyu 2003; Lejeune et al. 2003; Yamashita et al. 2005).

Importantly, the understanding of poly(A) tails, deadenylation, and decapping in normal (non-PTC-containing) mRNA metabolism improved over the last several years (reviewed in (Zhang et al. 2023)). Contemporary studies suggest that deadenylation is a part of normal mRNA maturation as well as normal mRNA decay (Chang et al. 2014; Lima et al. 2017; Yi et al. 2018; Eisen et al. 2020; Alles et al. 2023; Park et al. 2023; Tudek et al. 2021). Nascent mRNAs have a long poly(A) tail that is partially deadenylated to a ∼30-70 poly(A) tail as the mRNA matures. Another set of reactions can shorten the poly(A) tail and expose the 3’ end of the message, facilitating decapping at the 5’end and exonucleolytic degradation of the mRNA. How these two periods of deadenylation relate is unclear, but existing data support both occurring widely. Normal mRNAs’ stability correlates well with deadenylation rates in mouse fibroblasts, but NMD targets are notable exceptions, suggesting NMD targets are degraded independent of deadenylation and/or decapping (Eisen et al. 2020). This conflicts with the extant literature on NMD, confounding a mechanistic understanding of mRNA attack during NMD.

Several technical challenges complicate analysis of the effects of deadenylation, endonucleolytic cleavage, and exonucleolytic decay during NMD. A common technique to study NMD is short-read RNA-sequencing. Short read sequencing provides ensemble measures of fragments of mRNAs produced from a gene, which combined with the short nature of reads, prevents mRNA isoform assignment for many reads. Most short read RNA-seq techniques lose poly(A) tail information. For short read RNA sequencing techniques that retain poly(A) tail information, it is difficult or impossible to assign most poly(A) tail-containing reads to NMD isoforms as the PTC-introducing event is typically hundreds (or thousands) of bases away from the poly(A) tail. Short read sequencing techniques also require RNA fragmentation which loses endogenous cleavage information. Additionally, short read sequencing requires PCR; PCR of the homopolymeric poly(A) tail is notoriously inefficient, error-prone, and bias-inducing, compromising poly(A) tail information. While gel-based (*e*.*g*., northern blots) circumvent some of these issues, they are also low-throughput, yield comparatively little sequence information, and are still ensemble-based measurements of gene expression. However, recently developed long-read, whole-transcript, single-molecule mRNA sequencing techniques promise to circumvent these issues (for a more complete discussion of short and long-read sequencing, see (Hu et al. 2021)).

We aimed to determine the role of deadenylation in NMD and its relationship to endonucleolytic cleavage via nanopore direct RNA-sequencing. Our approach provides isoform, cleavage site, and poly(A) tail information for single, native mRNA molecules as they exist *in vivo*. Our analysis supports the idea that NMD targets are deadenylated during their maturation but not as a part of PTC-mediated degradation. By capturing mRNAs from NMD factor mutant backgrounds, we show that both SMG-6 and SMG-5 are required for endonucleolytic cleavage of a common set of NMD targets. Our work supports the idea that there is a single endonucleolytic cleavage reaction elicited by SMG-5/6 that is essential to NMD. In light of prior work on deadenylation and SMG-5, our results represent an important step forward in delineating NMD mechanism.

## RESULTS

### Nanopore degradome sequencing captures full-length mRNAs and degradation intermediates

To study NMD, we captured full-length and cleaved mRNAs using a degradome sequencing protocol (5TERA-seq, (Ibrahim et al. 2021)) (Fig 1A). (Protocol described in Methods.) Briefly, to capture mRNAs and enable nanopore sequencing, we annealed and ligated an ONT (Oxford Nanopore Technologies) 3’ adapter using an oligo-dT splint. The oligo-dT splint requires a minimum of 10 adenosines on the 3’end of an RNA molecule; poly(A) tails are >25 adenosines in length (Eisen et al. 2020; Lima et al. 2017). We performed no additional poly(A) selection as poly(A) selection can irreproducibly skew the view of the transcriptome (Viscardi and Arribere 2022). To differentiate cleaved RNAs, we subjected total RNA samples to 5’ adapter ligation using an end chemistry-based ligation specific to RNAs with 5’ monophosphates that arise from endonucleolytic cleavage or decapping (Ibrahim et al. 2021). The strategy is similar to other degradome sequencing methods (Ottens et al. 2017; Schmidt et al. 2015; Won et al. 2020), with the following important differences: (1) we performed full-length RNA sequencing without PCR, circumventing PCR-dependent biases in molecule capture, (2) the 3’ ONT adapter is added at the end of the poly(A) tail, thus retaining information on the entirety of the RNA molecule from its poly(A) tail through its 5’ end, and (3) the library contains both full-length and cleaved mRNAs, with the latter identified *in silico* via the presence of the 5’ adapter. To enhance the stability of cleaved mRNAs, we also performed a knockdown of the primary 5’ to 3’ exonuclease *xrn-1* via RNAi (Fig 1B). Overall, reads with the 5’ adapter were low abundance (∼2-5% of libraries; Table S1), as expected, since degradation intermediates are transient species and a minority of the overall RNA pool.

**Figure 1:**
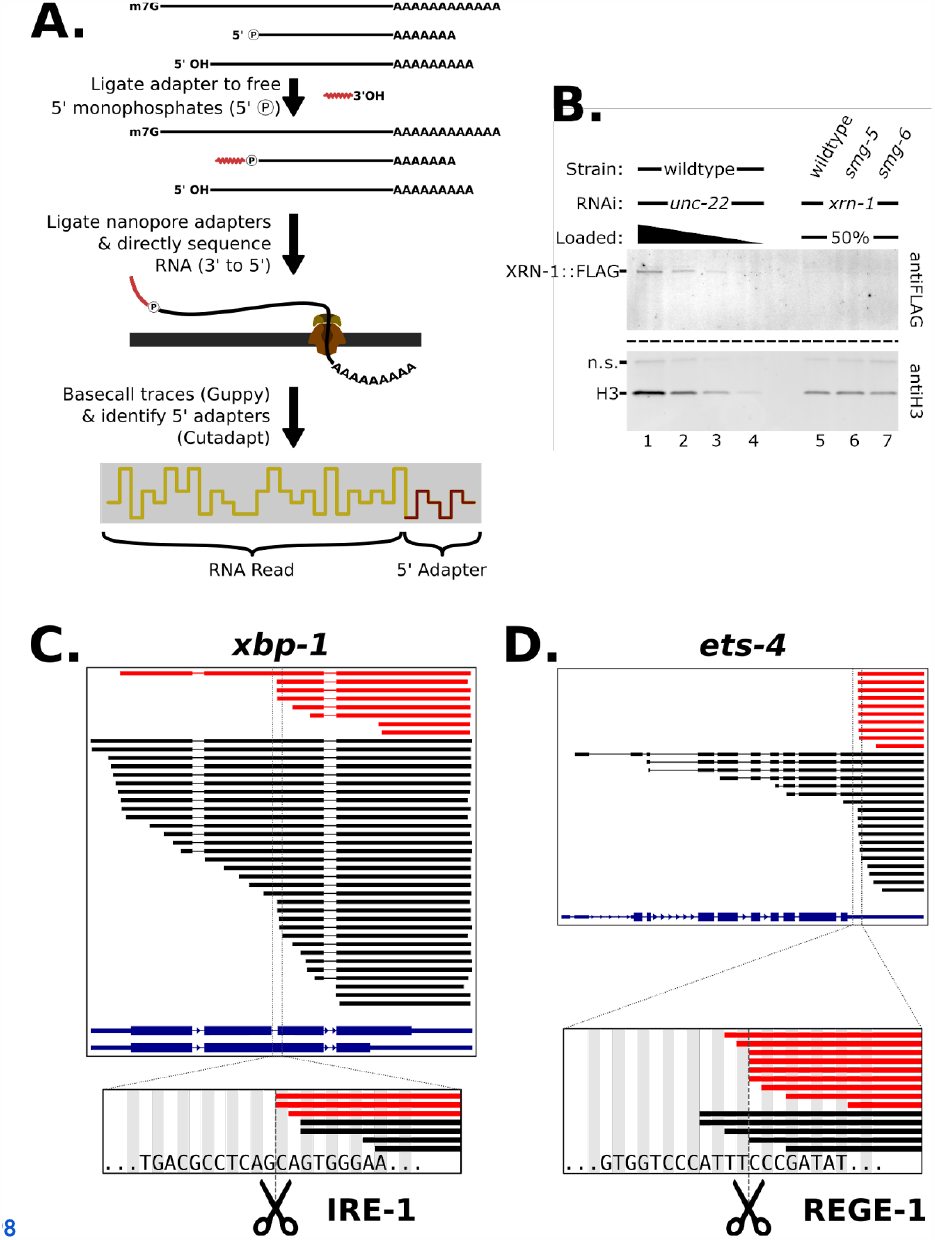
Workflow for the capture of mRNA degradation intermediates. A. Strategy to identify RNAs with a 5’ monophosphate using Oxford Nanopore Technologies’ direct RNA sequencing. Further detailed in methods. B. Western blot of XRN-1 protein knockdown by RNAi. C. (and D) 5’ monophosphate ends left on known endonuclease targets *xbp-1* (and *ets-4*). (Top) Red lines indicate molecules that contained the 5’ adapter sequence, and were thus derived from 5’ monophosphate-containing RNAs. Black lines indicate molecules that did not contain the 5’ adapter sequence. Thick lines indicate aligned sections of the RNA sequences and thin lines indicate alignment gaps of RNA reads that span annotated introns. The isoform annotations for *xbp-1* (and *ets-4*) are indicated in blue. The thickest sections of the isoform annotations indicate the coding sequences. (Bottom) A zoomed window showing reads with 5’ ends near the known endonuclease cleavage sequence and site (indicated with dotted line and scissors).

We performed nanopore direct RNA degradome sequencing with *C. elegans* total RNA and validated the technique’s performance using known endogenous endonuclease targets: *xbp-1* (cut by IRE-1) and *ets-4* (cut by REGE-1) (Habacher et al. 2016; Shen et al. 2001; Arribere and Fire 2018; Kim et al. 2022). For both *ets-4* and *xbp-1*, we observed adapted 5’ ends at and downstream of the previously identified cleavage site. The assay provided a high degree of specificity as some RNA molecules had 5’ ends at the known cleavage site with single-nucleotide precision (Fig 1C, D). We also observed full-length molecules spanning the cleavage site, as expected from the recovery of uncleaved mRNAs. The “full-length” (*i*.*e*., unadapted) population includes some shorter mRNAs resulting from incomplete adapter ligation, *in vitro* RNA hydrolysis, and/or pore sequencing dropoff. For simplicity, we describe RNAs as belonging to “cleaved” or “full-length” populations based on the presence or absence of the 5’ adapter, respectively.

Visual inspection of the library identified abundant cleavages within several additional endogenous genes, including *rps-22, rpl-30*, and *ubl-1*. Each of these genes contains an mRNA isoform encoding a premature stop codon (PTC), suggesting a relation to NMD. Indeed, many ribosomal protein genes produce NMD-eliciting splicing isoforms, as identified in a prior study of endogenous *C. elegans* NMD targets (Mitrovich and Anderson 2000). In the cases of *rps-22* and *rpl-30*, the splicing event that creates the PTC enabled the unique assignment of full-length and cleaved mRNA molecules to either the NMD-isoform or the non-NMD isoform. By assigning reads to isoforms, we observed that the vast majority of cleaved mRNAs derived from *rps-22* and *rpl-30* were made from the PTC-containing isoform (Fig 2A-C). Conversely, most full-length mRNA molecules corresponded to the non-PTC-containing isoform. In the case of *ubl-1*, unique isoform assignment is possible for full-length molecules but not mRNAs with 5’ ends downstream of the PTC due to a lack of unique splice information post-cleavage; we classify such molecules as “ambiguous”.

**Figure 2:**
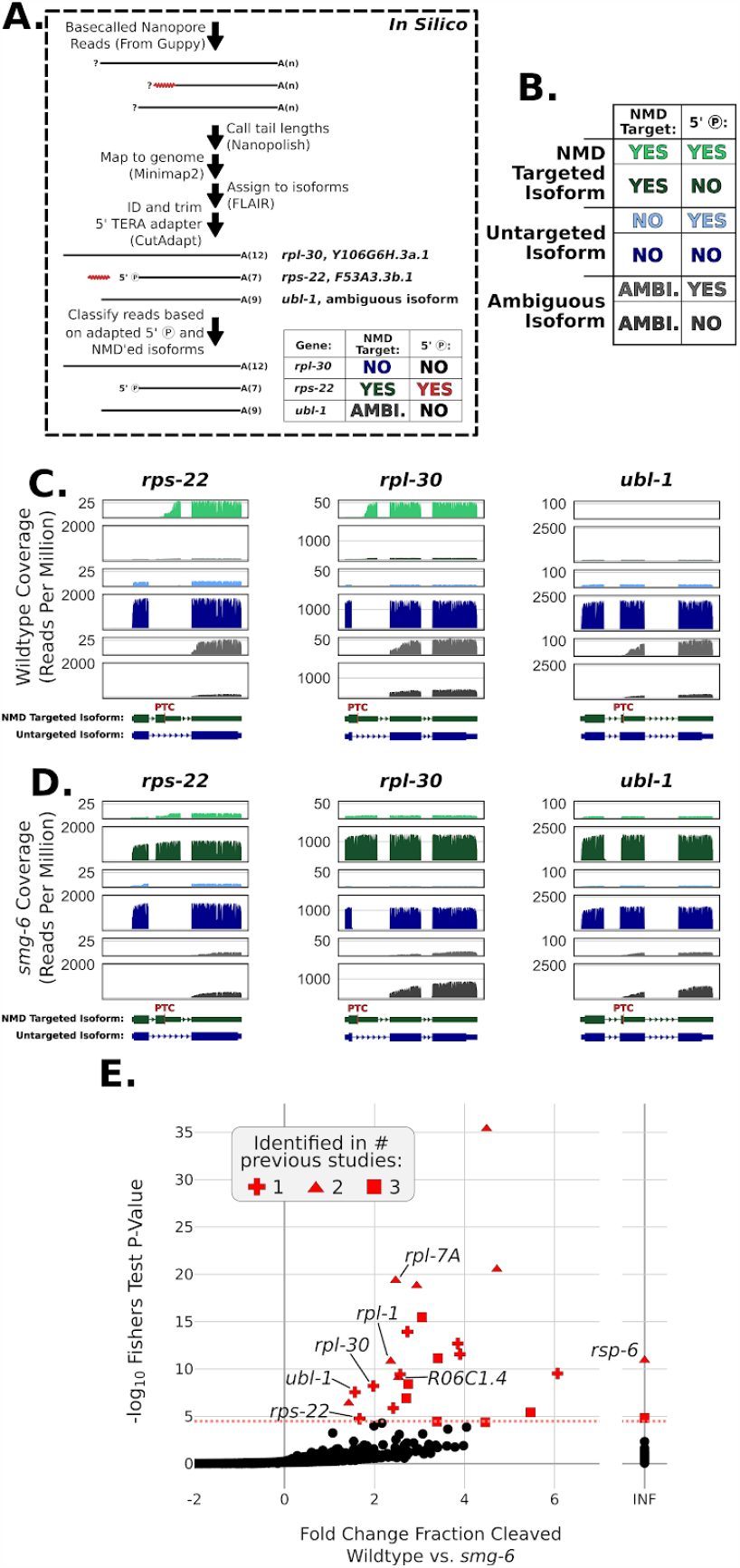
Nanopore degradome sequencing captures *smg-6*-dependent degradation intermediates on NMD targets. *A. In silico* read processing workflow for degradome sequencing. Details in methods. B. Key, indicating colors and order of isoform identity and 5’ monophosphate presence. C. Coverage plots of NMD targets’ (*rps-22, rpl-30*, and *ubl-1*) loci in wildtype animals. Read coverage (y-axes) are shown in normalized reads per million. From top to bottom, coverages are for the following categories: cleaved NMD isoforms (light green), full-length NMD isoforms (dark green), cleaved non-NMD isoforms (light blue), full-length non-NMD isoforms (dark blue), cleaved ambiguous isoforms (light gray), and full-length ambiguous isoforms (dark gray). Annotations at the bottom indicate the NMD-targeted isoforms and the primary non-NMD-targeted isoforms. The “PTC” indicates the location of the NMD-eliciting stop codon. D. Coverage plots of NMD targets’ loci in *smg-6* mutant animals. *E. De novo* identified NMD targets. Fisher’s Exact test was used to compare a contingency table of cleaved and full-length counts between wildtype and *smg-6* animals. Only genes containing 100 cumulative reads between the wildtype and *smg-6* libraries were used for this analysis. X-axis is the Log2 fold change between the fraction of cleaved reads (cleaved read count / total read count) for wildtype and *smg-6* animals. The salmon dashed line indicates the Bonferroni corrected P-value cutoff of 4.227E-5. For all genes above the Bonferroni corrected P-value cutoff, shapes indicate the number of previous studies that identified the gene; see also Table S2. “INF” indicates an infinite fold-change, caused by the complete absence of cleaved reads in *smg-6*.

The low abundance of full-length PTC-containing isoforms (*rps-22, rpl-30*, and *ubl-1*) alongside the abundance of cleaved PTC-containing isoforms (*rps-22* and *rpl-30*) demonstrates that our strategy captures endogenous NMD processes, with an abundant NMD intermediate being mRNAs cleaved downstream of the PTC.

### Degradome sequencing captures *smg-6*-dependent degradation intermediates on NMD targets

To understand the requirements for mRNA cleavage during NMD, we first analyzed the relationship of cleaved PTC-containing mRNAs to SMG-6. SMG-6 is noteworthy because it contains a PIN nuclease domain required for NMD and mRNA cleavage (Gatfield et al. 2003; Glavan et al. 2006; Eberle et al. 2009). We performed degradome sequencing of RNA isolated from animals carrying a previously validated mutant of an active site residue on SMG-6 (Kim and Modena et al. 2022), and determined isoform identity and cleavage status of RNA molecules. While abundance and distribution of non-PTC isoforms of *rps-22, rpl-30*, and *ubl-1* were similar in the *smg-6* mutant (Fig 2D, dark blue), PTC-containing isoforms were affected in two key ways: (1) the *smg-6* mutant exhibited loss of mRNA fragments cleaved at and downstream of the PTC (Fig 2, compare light green in C and D), and (2) the *smg-6* mutant exhibited high levels of full-length mRNAs on the PTC-containing isoform (Fig 2, compare dark green in C and D). These observations are consistent with a model in which the nuclease activity of SMG-6 is required for PTC-proximal mRNA cleavage.

Among cleaved mRNAs, we also noted a minor population with 5’ends mapping near the 5’end of both normal and NMD targeted isoforms in wildtype and *smg-6* animals (Fig 2C, D, light green and light blue), likely the product of mRNA decapping. The low abundance of uncapped mRNAs on NMD targets compared to downstream, PTC-proximal cleaved mRNAs suggests that decapping is a relatively minor contributor to NMD target decay in this system (see Discussion).

### Analysis of *smg-6*-dependent degradation intermediates identifies primary NMD targets *de novo*

The abundance of degradation intermediates in wild-type animals and their rarity in *smg-6* animals provides a means to identify direct NMD targets *de novo*. We quantified cleaved and uncleaved molecules in wild-type and *smg-6* animals in a 2x2 contingency table and then applied Fisher’s Exact Test with a Bonferroni-corrected P-value cutoff to identify genes for which the frequency of cleaved mRNAs decreased in *smg-6* animals (Fig 2E, see a complete description in Methods).

Of 1,183 genes that passed cutoffs for consideration in the analysis, 25 exhibited a significant reduction in cleaved mRNAs in the *smg-6* mutant. All 25 genes (including *rpl-30, rps-22*, and *ubl-1*) were previously identified by at least one prior NMD study in *C. elegans* (Table S2) (Muir et al. 2018; Kim et al. 2022; Mitrovich and Anderson 2000; Ramani et al. 2009). Visual analysis of these 25 genes revealed that 23 contained a obvious NMD-eliciting feature, such as an upstream Open Reading Frame (uORF; *smd-1, rpl-1, C45B2*.*8, farl-11*, and *zip-12*) or a 3’ Untranslated Region (3’UTR)-contained intron (*Y73B3A*.*18, rpl-7A, rpl-30, F19B2*.*5, nhr-114, C30E1*.*9, rsp-6, tos-1, ubl-1, C53H9*.*2, rpl-3, T05E12*.*6, H28G03*.*2, C35B1*.*2, ddo-2, R06C1*.*4*, and *rps-22*). The remaining 2 genes (*col-182* and *Y39B6A*.*21*) lacked an obvious NMD-inducing feature, though contained a *smg-6*-dependent accumulation of cleaved reads near the stop codon, suggesting that the stop codon of these mRNAs is targeted as a PTC through yet unknown mechanisms. Thus, statistical analysis of degradation fragment abundance across wild-type and NMD-deficient strains can identify NMD targets *de novo*.

We expect that the fraction of NMD targets identified in this analysis (25 of 1,183 genes, ∼2.1%) represents a conservative, lower bound on the overall frequency of NMD targets. Indeed, among a list of abundant NMD targets (Mitrovich and Anderson 2000), we did not identify *rpl-12* as a target. Visual inspection revealed that the PTC-containing isoform of *rpl-12* was expressed as a low fraction of all *rpl-12*-derived transcripts. Failure to identify *rpl-12* was thus expected as our statistical analysis was performed on read counts tabulated by gene rather than mRNA isoform. In principle, the approach could be extended to identify individual isoforms targeted by NMD, though, in practice, we found existing isoform annotations and isoform-assignment tools inadequate for the task (see Methods). As annotations, isoform-assignment tools, and sequencing depth continue to improve, this approach will identify additional NMD targets.

### NMD target mRNAs and their degradation products have poly(A) tails that are as long – or longer – than non-NMD target mRNAs

A significant advantage of a direct RNA-seq technique is our ability to capture information on a per-molecule basis. Among the information captured is an mRNA’s poly(A) tail length. The existing NMD literature suggests a contribution from deadenylation (Loh et al. 2013; Chen and Shyu 2003; Lejeune et al. 2003; Yamashita et al. 2005), and our nanopore libraries seemed well-suited to investigate this contribution. The categorization of molecules as PTC-containing (or not) and as cleaved (or not) allowed for a comparison of poly(A) tail lengths across these categories. Our degradome sequencing approach thus provides a novel way to assess the role of deadenylation and poly(A) tails in NMD.

Analysis of full-length mRNAs genome-wide revealed a distribution of poly(A) tail lengths with a median length of ∼52 adenosines (Fig 3A). This length distribution is consistent with others’ measurements of poly(A) tails in *C. elegans* via long-read sequencing (Roach et al. 2020; Legnini et al. 2019), short-read sequencing (Lima et al. 2017), and gels (Nousch et al. 2017). Focusing on cleaved mRNAs, we observed an overall distribution similar to full-length species (Fig 3A). Thus our libraries recovered poly(A) tail information in line with prior measurements.

**Figure 3:**
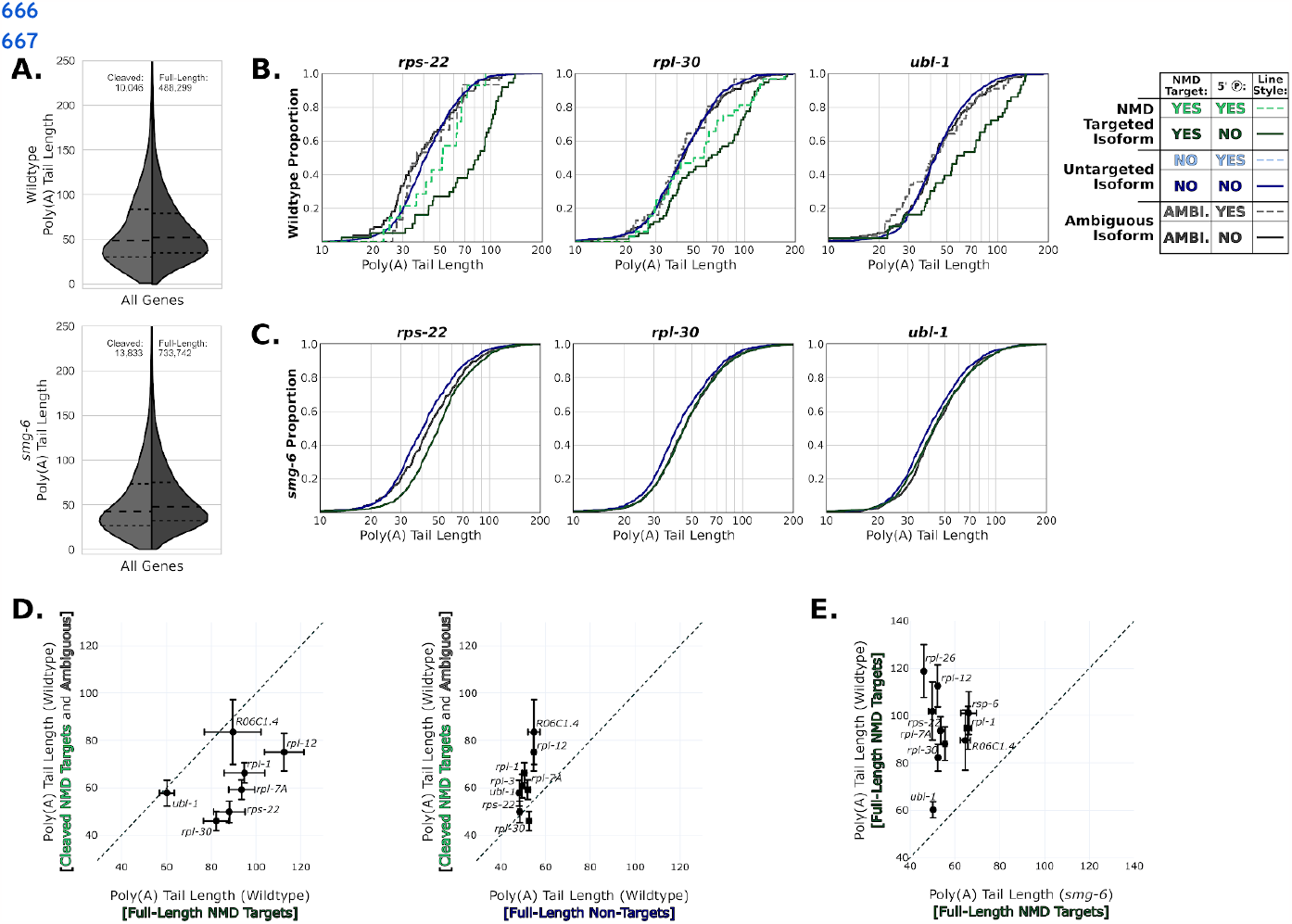
NMD target mRNAs and their degradation products have poly(A) tails that are as least as long as normal mRNAs. A. Violin plots for all mRNAs in wildtype and *smg-6* animals. The left side of each violin (in light gray) shows the distribution of cleaved reads’ tail lengths, while the right (in dark gray) shows full-length reads. Long dashed lines indicate the means and short dashed lines indicate 1st and 4th quartile boundaries. B. Poly(A) tail length cumulative distribution function (CDF) plots of example genes (*rps-22, rpl-30*, and *ubl-1*) in wild-type animals. The same color scheme is used here as in Figure 2: cleaved NMD isoforms (dashed light green), full-length NMD isoforms (dark green), cleaved non-NMD isoforms (dashed light blue), full-length non-NMD isoforms (dark blue), cleaved ambiguous isoforms (dashed light gray), and full-length ambiguous isoforms (dark gray). For each plot, only categories that had at least 10 poly(A) tail-called reads are shown. See also Table S3 for statistical analysis. C. Poly(A) tail length CDF plots of example genes in *smg-6* animals. Low counts of cleaved reads on these loci cause more cleaved, PTC-mapping RNA species to fall below the 10 read cutoff. See also Table S3 for statistical analysis. D. Comparisons of mean poly(A) tail lengths between cleaved versus full-length NMD targets and cleaved NMD targets versus full-length non-targets for genes in which we could identify the NMD target isoform. The dashed line indicates the diagonal where X = Y. Error bars indicate the standard error of the mean. Only genes with at least 10 reads in each category are shown. E. Similar to subfigure D: Comparison of poly(A) tail lengths between full-length NMD targets in wildtype animals versus *smg-6* animals.

To better understand the relationship between poly(A) tails and NMD, we focused on mRNA molecules produced from the exemplary genes *rps-22, rpl-30*, and *ubl-1* (Fig 3B). Cleaved mRNAs from these genes exhibited poly(A) tail lengths similar to the genome-wide distribution and similar to full-length normal mRNAs produced from the same genes. Thus, at (or immediately after) the time that cleavage occurs, NMD targets have a poly(A) tail length similar to that of non-NMD targets. Put another way, the existence of polyadenylated, cleaved NMD targets is inconsistent with complete deadenylation as a prerequisite for cleavage.

To gain further insight into the life cycle of poly(A) tails during NMD, we examined full-length NMD target mRNAs (Fig 3B). Full-length NMD target mRNAs would be expected to include mRNAs that have not yet entered the translational pool and are yet to be targeted by NMD (*e*.*g*., recently transcribed mRNAs). Full-length PTC-containing mRNAs exhibited significantly longer tails than either cleaved, PTC-containing mRNAs or full-length, non-PTC-containing mRNAs. The effect was seen for PTC-containing isoforms in each of *rps-22, rpl-30*, and *ubl-1* (Fig 3B, accompanying statistical analysis Table S3) as well as each of the other 14 genes where we could identify the NMD-eliciting isoform (Fig S1). The reduction in poly(A) tail lengths between full-length NMD target mRNAs and cleaved NMD target mRNAs supports NMD targets experiencing deadenylation prior to or coincident with endonucleolytic cleavage.

Thus, by the time that they experience cleavage, NMD targets’ poly(A) tails are similar in length (or longer than) normal mRNAs’ poly(A) tails.

### Upon knockout of *smg-6*, NMD target mRNAs have similar poly(A) tail lengths to normal mRNAs

We considered two models to explain NMD target mRNAs’ long poly(A) tails in wild-type animals:

1. Maturation-dependent deadenylation: full-length NMD targets’ long poly(A) tails are due to their relative youth among cellular mRNAs. As NMD targets are targeted for degradation early in their life span, a higher proportion of full-length NMD target mRNAs will be nascent. As nascent mRNAs have longer poly(A) tails than mature mRNAs, NMD targets’ poly(A) tails will be longer on average. In this model, the increased poly(A) tail lengths of full-length NMD targets are a side-effect of the absence of a stable, mature mRNA population.
2. NMD-dependent deadenylation: NMD targets experience deadenylation during their decay. In this model, NMD target mRNAs have longer poly(A) tails, and in the course of NMD, their poly(A) tails are shortened towards a length that resembles normal mRNAs. In this model, deadenylation is central to the decay reaction(s) of NMD.

As an initial test between these models, we examined mRNA molecules in *smg-6* mutant animals. In *smg-6* animals, NMD targets are not cleaved, and full-length mRNA targets accumulate (Fig 2D,E). The maturation-dependent deadenylation model predicts that poly(A) tails of NMD targets will resemble those of normal mRNAs in an NMD mutant background. In contrast, the NMD-dependent deadenylation model predicts that NMD targets will maintain their longer poly(A) tails due to the loss of NMD and associated deadenylation that accompanies NMD loss. Our results show that the poly(A) tails of NMD target mRNAs are significantly shorter in *smg-6* animals relative to wild-type animals and are similar to that of normal mRNAs (Fig 3C-E, Fig S1). This result is consistent with the maturation-dependent deadenylation model.

### NMD target poly(A) tail lengths resemble non-NMD targets upon *smg-5* knockout

The existing literature suggests that the heterodimer SMG-5/7 promotes deadenylation during NMD. Because this role for SMG-5/7 would fit into the NMD-dependent deadenylation model, we entertained a variation to this model that could fit the data shown thus far: SMG-5/7 deadenylate NMD targets before cleavage via SMG-6. In this model, full-length NMD targets would accumulate with a shorter poly(A) tail in *smg-6* animals compared to wild-type, as is the case (Fig 3C,E). To distinguish between the maturation-dependent and the revised NMD-dependent deadenylation models, we considered the expectations of a *smg-5* mutant. Under the revised model, we would expect full-length NMD targets to accumulate in *smg-5* animals with long poly(A) tails. However, if the maturation-depended deadenylation model were correct, in *smg-5* animals we would again expect NMD targets to accumulate with normal-length poly(A) tails.

We performed degradome sequencing in a *smg-5* mutant and analyzed poly(A) tail lengths. In *smg-5* animals, normal mRNAs’ poly(A) tails were comparable to wild-type (Fig 4A,B). However, full-length NMD targets’ poly(A) tails were notably shorter than in wild-type, and were similar to normal mRNAs’ poly(A) tail lengths (Fig 4B,C, Fig S1). This result is consistent with results in *smg-6* animals (Fig 3C, 4D, S1) and the maturation-dependent deadenylation model, but would not be expected under the NMD-dependent deadenylation model.

**Figure 4:**
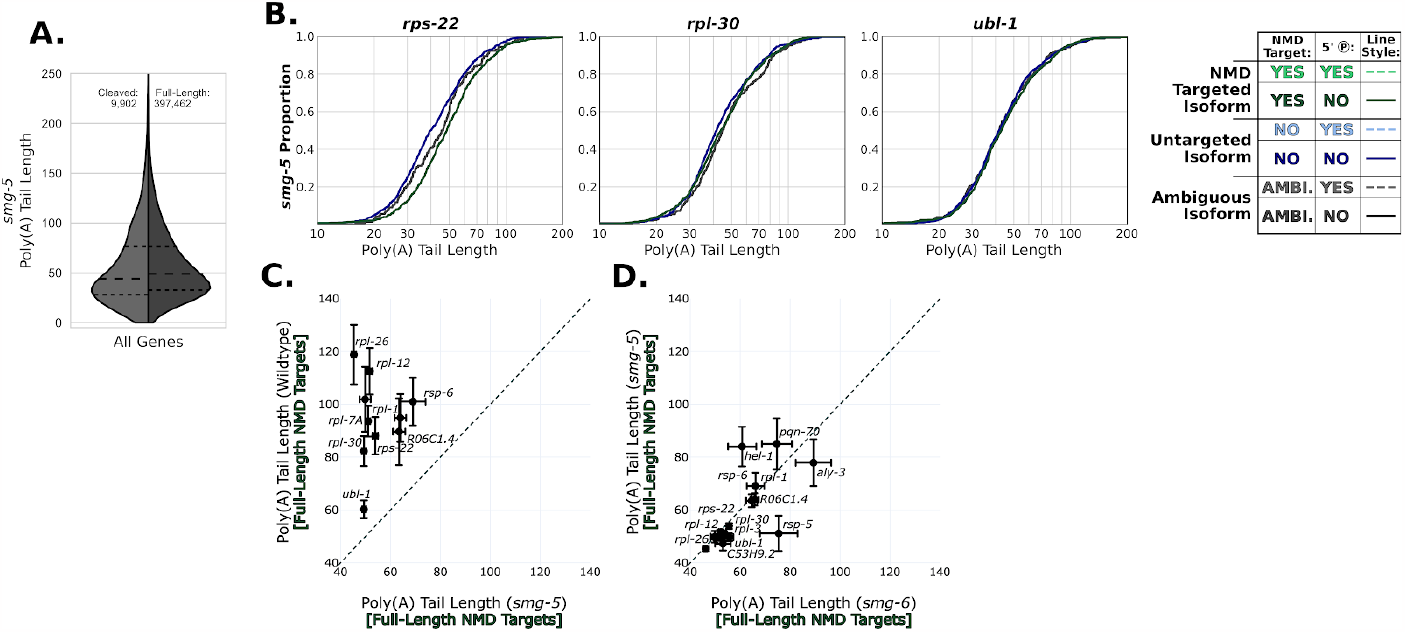
NMD Target poly(A) tail lengths resemble normal mRNAs in *smg-5* animals. A. Violin plots for all mRNAs in *smg-5* animals. The left side of the violin (in light gray) shows the distribution of cleaved reads’ tail lengths, while the right (in dark gray) shows uncleaved reads. Long dashed lines indicate the means and short dashed lines indicate 1st and 4th quartile boundaries. B. Poly(A) tail length cumulative distribution function (CDF) plots of example genes (*rps-22, rpl-30*, and *ubl-1*) in wild-type animals. The same color scheme is used here as in Figures 2 and 3: cleaved NMD isoforms (dashed light green), full-length NMD isoforms (dark green), cleaved non-NMD isoforms (dashed light blue), full-length non-NMD isoforms (dark blue), cleaved ambiguous isoforms (dashed light gray), and full-length ambiguous isoforms (dark gray). For each plot, only categories that had at least 10 poly(A) tail-called reads are shown. See also Table S3 for statistical analysis. C. Comparison of poly(A) tail lengths between full-length NMD targets in wildtype animals versus *smg-5* animals. The dashed line indicates the diagonal where X = Y. Error bars indicate the standard error of the mean. Only genes with at least 10 reads in each category are shown. D. imilar to subfigure C. Comparison of poly(A) tail lengths between full-length NMD targets in *smg-6* animals versus *smg-5* animals. A larger number of genes are shown here due to more genes passing the cutoff of 10 reads in each category.

Our analysis of poly(A) tail lengths in wild-type, *smg-6*, and *smg-5* animals is consistent with the idea that the extent of deadenylation experienced by NMD targets is within the scope of that experienced during normal mRNA maturation. In the Discussion, we expound on this idea and relate it to the extant literature on deadenylation and NMD.

### *smg-5* is required for *smg-6*-dependent cleavages on NMD targets

The lack of a clear NMD-dependent deadenylation signature suggested that SMG-5’s hypothesized role in deadenylation merited revision. It is well-established that SMG-5 is required for NMD across animal systems (Hodgkin et al. 1989; Nelson et al. 2018; Baird et al. 2018; Alexandrov et al. 2017; Zhu et al. 2020; Zinshteyn et al. 2021; Huth et al. 2022). We therefore examined the *smg-5* animal transcriptome for clues as to SMG-5’s biochemical function(s).

Upon knockout of *smg-5*, we noted a loss of degradation intermediates on NMD-sensitive isoforms, similar to the loss seen in *smg-6* animals (Fig 5A, Fig 2D). The effect was specific to NMD targets, as targets of other endonucleases (*ets-4* and *xbp-1*) were unaffected in the mutants (Fig S2). The loss of cleavage fragments in *smg-5* and *smg-6* animals is consistent with the idea that both proteins collaborate in a single pathway to cleave a common set of mRNAs. To ascertain the extent to which SMG-5 and SMG-6 impact a similar set of targets, we performed *de novo* target identification in *smg-5* animals (Fig 5B, as described in Fig 2E). All 15 of 15 *smg-5* targets were identified as *smg-6* targets (Fig 5C, Table S2). The remaining 10 *smg-6*-specific targets fell just below the p-value or read count cutoffs in the *smg-5* analysis, a likely result of decreased depth in the *smg-5* degradome sequencing library (Table S1). Consistent with this, all 25 *smg-6* targets were also increased in the *smg-5* mutant.

**Figure 5:**
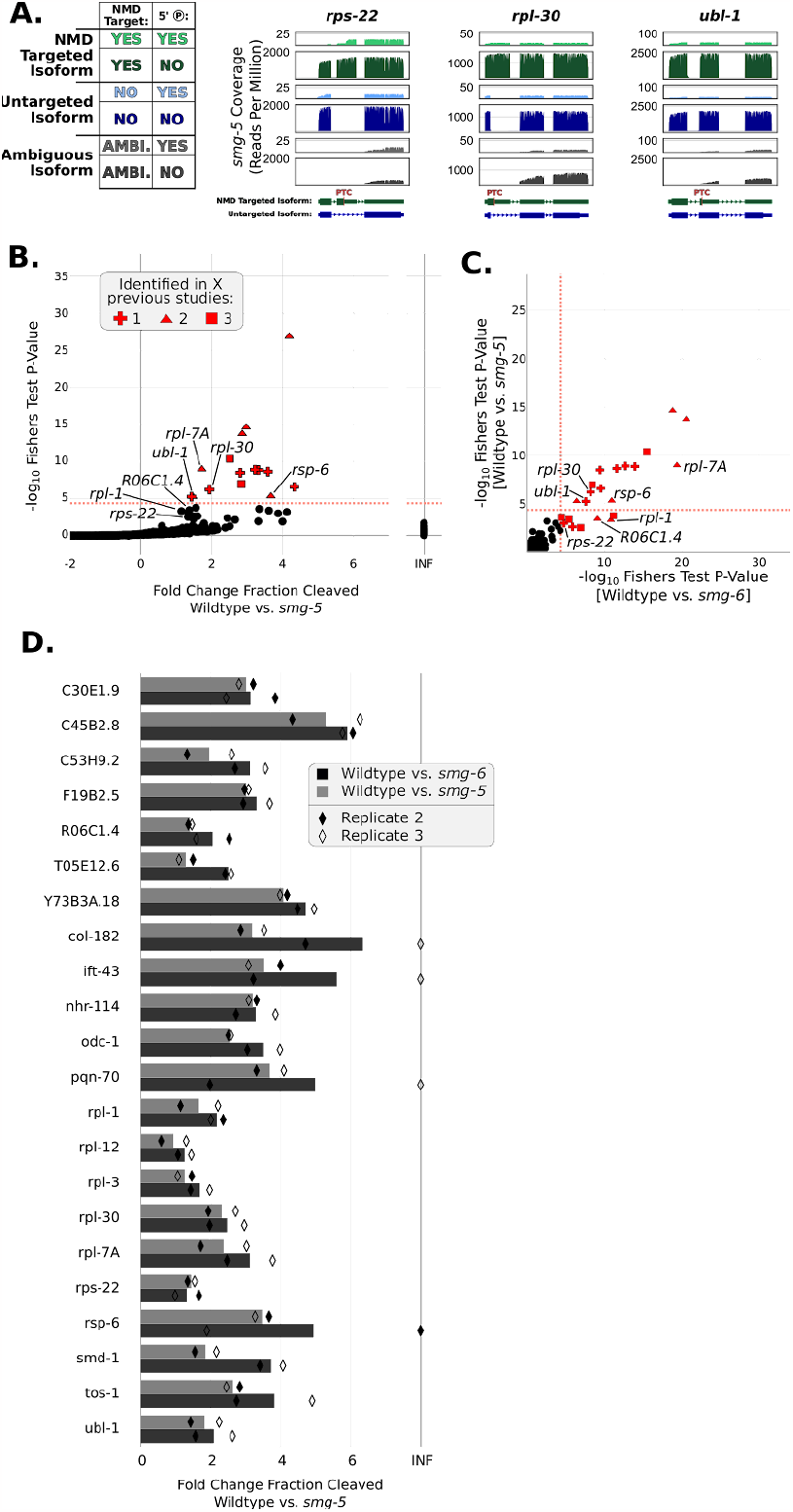
*smg-5* is required for *smg-6*-dependent cleavages on NMD targets. A. Coverage plots of NMD targets’ (*rps-22, rpl-30*, and *ubl-1*) loci in *smg-5* animals. Read coverage (y-axes) are shown in reads per million. From top to bottom, coverages are for the following categories: cleaved NMD isoforms (light green), full-length NMD isoforms (dark green), cleaved non-NMD isoforms (light blue), full-length non-NMD isoforms (dark blue), cleaved ambiguous isoforms (light gray), and full-length ambiguous isoforms (dark gray). Annotations at the bottom indicate the NMD-targeted isoforms and the primary non-NMD-targeted isoforms. The “PTC” indicates the location of the NMD-eliciting stop codon. B. *De novo* identified NMD targets in *smg-5*. Fisher’s Exact test was used to compare a contingency table of cleaved and full-length counts between wildtype and *smg-5* animals. Only genes containing 100 cumulative reads between the wildtype and *smg-5* libraries were used for this analysis. X-axis is the Log2 fold change between the fraction of cleaved reads (cleaved read count / total read count) for wildtype and *smg-5* animals. The salmon dashed line indicates the Bonferroni corrected P-value cutoff of 4.227E-5. For all genes above the Bonferroni corrected P-value cutoff, shapes indicate the number of previous studies that identified the gene (see also Table S2). C. Correlation of Fisher’s Exact test results for *de novo* NMD target analysis in *smg-5* and *smg-6* animals. The salmon dashed lines indicate the Bonferroni corrected P-value cutoff of 4.227E-5. Again, shapes indicate the number of previous studies that identified the gene as an NMD target. D. Bar plots of Log_2_ fold change between the fraction of cleaved reads (cleaved read count / total read count) for wildtype and *smg-5* animals or wildtype and *smg-6* animals among targets identified by Fisher’s Exact test analysis.

A biological replicate of these experiments reproduced our observations (Fig S1, S3-S5). Taken together, these results support a model in which SMG-5 and SMG-6 act in a single pathway to cleave NMD targets.

## DISCUSSION

The data we present are consistent with a model with the following steps of NMD in animals: (1) like normal (PTC-lacking) mRNAs, nascent, PTC-containing mRNAs are produced with long and heterogeneous poly(A) tails; (2) like normal (PTC-lacking) mRNAs, around the onset of translation, PTC-containing mRNAs’ poly(A) tails are shortened to a length similar to mature mRNAs; (3) after the onset of translation, PTC-containing mRNAs are targeted by NMD machinery and cut near their PTC in a reaction that requires SMG-5 and SMG-6; (4) the downstream cleavage fragment is then cleared via XRN-1. We propose that NMD targets’ deadenylation is part of mRNA maturation rather than a consequence of decay.

This model is consistent with existing data on deadenylation of normal mRNAs. Normal mRNAs emerge from the nucleus with long and heterogeneous poly(A) tails, and subsequently experience a rapid period of deadenylation (Eisen et al. 2020; Sawicki et al. 1977; Tudek et al. 2021). Normal mRNAs can also experience a more variable (and often much slower) deadenylation over their lifetime that is associated with translation (Eisen et al. 2020; Yi et al. 2018; Tudek et al. 2021; Park et al. 2023). NMD targets are only recognized as such during translation, predicting that PTC-containing nascent mRNAs will also experience the initial period of deadenylation. Since NMD targets are not found in the translationally stable pool of mRNAs, they do not experience the second period of deadenylation. While both periods of deadenylation are thought to occur after nuclear export, their precise relationship to translation and translation termination remains unclear. Future studies in this area will prove informative, both for an understanding of normal mRNA metabolism and the metabolism of PTC-containing mRNAs.

The model is also consistent with existing data on deadenylation during NMD, though our interpretation of prior data differs. Of three studies that directly examined poly(A) tails on NMD targets, each observed that NMD targets experience partial deadenylation to some non-zero poly(A) tail length prior to loss of the mRNA (for example, Fig 3B of (Yamashita et al. 2005); Fig 6B,C of (Lejeune et al. 2003); Fig 1C of (Chen and Shyu 2003)). At the time these studies were done, it was generally thought that longer poly(A) tails were associated with more stable mRNAs, and so the deadenylation seen on NMD targets was invoked as a mechanism of NMD. Knowing now that mRNAs are born with long poly(A) tails that are subsequently shortened, the deadenylation signature in these studies may merely be a consequence of nascent mRNA maturation. Deadenylation appears heightened (and is more easily visualized) on NMD targets because of the lack of a stable, translationally mature, partially deadenylated mRNA population. Thus it is possible that deadenylation on NMD targets is not directly causal in NMD target mRNAs’ accelerated decay. Again, a better understanding of the deadenylation experienced by nascent mRNAs will reveal the extent to which deadenylation is similar–or differs–between normal mRNAs and NMD targets.

PTC-associated 5’monophosphate ends are in principle consistent with either of SMG-6-mediated endonucleolytic cleavage or decapping followed by 5’>3’ exonuclease degradation, but we favor the former. In prior work (Kim et al. 2022), we captured RNA 3’ends at and upstream of the stop codon, a result expected from endonucleolytic cleavage near the PTC but not expected from decapping. We note a requirement for at least ten 3’ terminal adenosines for capture with our nanopore strategy, preventing detection of RNAs with poly(A) tails shorter than ten or RNAs with non-adenosine terminal residues. Untemplated uracils can occur on mRNAs with poly(A) tails <25 adenosines (Eisen et al. 2020). The capture of such RNAs (and their relationship to NMD) will require protocols different from those used here.

Interestingly, we noticed that some NMD targets exhibited slightly longer poly(A) tails even in NMD mutants, suggesting that PTC-containing mRNAs are mildly defective in deadenylation. For example, *rps-22* PTC-containing mRNAs exhibited longer poly(A) tails compared to *rps-22* non-PTC-containing mRNAs in wildtype (Fig 3B), *smg-6* (Fig 3C), and *smg-5* (Fig 4B). While the magnitude of the difference was smaller in *smg* animals (a median difference of ∼10 adenosines compared to ∼45 adenosines in wildtype), the effect was statistically significant. Thus, even in the absence of NMD, NMD targets experience weaker deadenylation compared to non-targets. This result may prove mechanistically informative for the understanding of deadenylation as it relates to translation; perhaps PTCs alter ribosome dynamics in such a way that NMD targets inefficiently recruit deadenylases compared to normal mRNAs.

Our results support a role for SMG-5 in endonucleolytic cleavage of NMD targets. Here, loss of SMG-5 or SMG-6 prevented PTC-proximal cleavages on a common set of NMD targets, suggesting that both factors collaborate to cut NMD targets. This is consistent with the identification of SMG proteins in *C. elegans*, which showed that loss of either SMG-5 or SMG-6 exhibited a similar de-repression of a common set of NMD targets (Hodgkin et al. 1989), and our work here suggests that SMG-5/6 do so via mRNA cleavage. While the picture is still solidifying in human cells, we note that the idea that SMG-5/6 work together in a single NMD pathway is supported by recent work (Boehm et al. 2021). A better understanding of the cleavage reaction will clarify how SMG-5 and SMG-6 collaborate to cut mRNAs.

A current limitation of the work is that we only have data on the ∼1,183 most highly expressed genes, among which we confidently identified a few dozen NMD targets. It is possible that the mechanisms uncovered here may vary among more lowly expressed genes. The models and techniques outlined here should make it possible to carry out such experiments–either via targeted long-read sequencing approaches or experiments that otherwise distinguish between cleaved and uncleaved NMD targets.

Here we showcase the ability to attain novel mechanistic insight into mRNA metabolism from long-read sequencing technology. Sequencing entire mRNA molecules from poly(A) tail through isoform/splicing information and 5’end/cleavage status allowed us to deconvolve properties of different mRNA populations that are difficult–or impossible–to study with other approaches. Coupled with the genetic tractability of *C. elegans*, this approach yielded novel insight into the NMD pathway. We expect similar approaches to be informative for various mRNA metabolism pathways across animals.

## ACKNOWLEDGEMENTS

We thank the Arribere Lab for feedback, Melissa Jurica, Manny Ares, Fadia Ibrahim, and Robert Hogg for comments on the manuscript. This work was supported by a T32 training grant (5T32GM133391) to M.J.V., Searle Scholars Award to J.A.A., and an R01 grant (R01GM131012) to J.A.A.

## DATA AVAILABILITY

Sequencing data is deposited at NCBI under SRA BioProject: PRJNA1010807.

## METHODS

### Strains

All *C. elegans* strains were made in N2 background animals (VC2010 (Thompson et al. 2013)) and a list of strains is available (Table S4).

### RNA interference knockdown of *xrn-1* and sample collection

All animals were grown under RNAi knockdown conditions of the primary 5’ to 3’ endonuclease, *xrn-1*, based on the methodology described in (Pule et al. 2019). Briefly, an RNAi feeding strain was made via transformation of an *xrn-1* targeting RNAi plasmid from (Kamath and Ahringer 2003) into *ht115(de3)*. The bacteria were grown overnight at 37C to an optical density of 4.0 in 2×YT medium containing carbenicillin (25 μg/mL). IPTG was added to a concentration of 100 μg/mL to induce dsRNA expression, and growth was continued for 4 hours. The feeding strain was then spread on NGM plates containing carbenicillin (25 μg/mL) and IPTG (100 μg/mL). The RNAi-expressing lawns were grown overnight at room temperature prior to the addition of *C. elegans* eggs (next).

Animals were bleached to obtain a synchronous population of eggs and then grown at 20C until L3/L4 stage on the above-described RNAi plates. Animals were collected on a sucrose cushion (5%) to minimize bacterial contamination, pelleted, and washed with EN50 and M9. Pellets were resuspended in TRIzol reagent (Ambion, cat#15596026), lysed by freeze-cracking, and total RNA was isolated by chloroform extraction. Total RNA integrity was assessed with the Agilent high-sensitivity RNA system for TapeStation. Only total RNA samples with RNA integrity number equivalent (RINe) values greater than 7.0 were used for sequencing.

### 5’ adapter ligation, and direct RNA sequencing

To identify RNAs containing 5’ monophosphates, we used 5TERA (Ibrahim et al. 2021) with modifications. Briefly, total RNA was subjected to a T4 RNA ligase reaction with an RNA adapter (JA-MV-25, /5Biosg/rArArUrGrArUrArCrGrGrCrGrArCrCrArCrCrGrArGrArUrCrUrArCrArCrUr CrUrUrUrCrCrCrUrArCrArCrGrArCrGrCrUrCrUrUrCrCrGrArNrNrN) at 37C for 3 hours. Ligated RNA was cleaned with Zymo Research RNA Clean & Concentrator kits (Cat#: R1019) using manufacturer specifications to remove RNA species smaller than 200nt, including adapter. Differing from the 5TERA protocol, we did not enrich for adapted RNAs.

For nanopore sequencing, 5 micrograms of ligated total RNA was used as input for Oxford Nanopore Technologies’ (ONT’s) direct RNA sequencing kit (dRNA-Seq, SQK-RNA002). dRNA-seq libraries were prepared according to the manufacturer’s instructions (protocol version as released: September 13th, 2021) with the following modification: (1) increased input, (2) use of total RNA as input, and (3) use of SuperScript IV (ThermoFisher Invitrogen, cat#18090010) rather than the ONT’s specification of SuperScript III (ThermoFisher Invitrogen, cat#18080051).

### Nanopore sequencing software, basecalling, and alignment

All raw voltage traces were collected as FAST5 files using Oxford Nanopore Technologies’ software MinKNOW (Core versions 4.3.4 to 5.3.1). In order to minimize variability in basecalling and downstream analyses based on software versions, all libraries were reprocessed from FAST5s with the same pipeline as follows. Raw FAST5 files from MinKNOW were basecalled with Guppy (v6.5.7+ca6d6af) in GPU mode using parameters: guppy_basecaller -c rna_r9.4.1_70bps_hac.cfg. Basecalled reads were aligned to the *C. elegans* genome (WBCel235) using MiniMap2 (v2.17-r941) (Li 2018) with recommended settings for dRNA-seq: minimap2 -x splice -uf -k14. Additionally, the parameter –junc-bed was used with a bed genome annotation file to provide minimap2 with splice junction information.

### Additional post-processing for degradome sequencing

To identify reads containing the 5’ adapter sequence (derived from RNAs that had 5’ monophosphates and underwent ligation), we utilized CutAdapt (Martin 2011) as specified in (Ibrahim et al. 2021). Fastq comments containing information from CutAdapt were integrated into post-alignment BAM/SAM files using the pysam library (https://github.com/pysam-developers/pysam) as SAM tags.

Nanopolish (Workman et al. 2019) was used with default parameters to assess poly(A) tail lengths in all sequenced libraries. For plots utilizing mean tail lengths as a summary statistic, we restricted analyses to genes with 10 or more reads. Information from basecalling, adapter identification, alignment, gene assignment, and tail-length calling was consolidated into extended SAM format files as additional tags. Reads were required to have successful mapping and gene assignment in order to be used for downstream analysis.

### *De novo* endonuclease target identification

To identify NMD targets *de novo*, we quantified cleaved and uncleaved read counts in wild-type and *smg-6* (or *smg-5*) animals in a 2x2 contingency table. We then used Scipy’s implementation of Fisher’s Exact Test (Virtanen et al. 2020) with a Bonferroni-corrected P-value cutoff to identify genes where the frequency of cleaved mRNAs decreased in mutant animals. In order to ensure adequate statistical power, we also imposed cumulative read count cutoffs of 100 reads per gene.

An important limitation is that this analysis was performed at the gene level and will miss many examples where individual isoforms change but not substantially enough to produce a detectable effect at the gene level. While we attempted to run the analysis at the isoform level, issues with annotations and accurate/unambiguous isoform assignment limited success. We found that manual identification of the NMD-eliciting isoforms was necessary to identify PTC sites. This is due to errors in systematic annotation software that will (for example) annotate the longest possible coding sequence that could be produced by an mRNA, rather than what is actually translated, which is often much shorter, more 5’, and more likely to generate a PTC. Additionally, annotations are built from wildtype transcriptomes, and thus the unstable mRNA isoforms found in NMD mutant animals are often absent. For previously identified NMD targets, we produced our overlap table by relying on each publication’s reported target list (Table S2).

### Isoform-level tail and coverage plots

Due to the above-noted limitations, only a subset of NMD-targeted genes had sufficient information to distinguish between NMD-targeted and untargeted isoforms. We utilized this list of 17 genes (*rpl-30, rps-22, ubl-1, rpl-7A, rpl-3, rpl-1, rpl-12, hel-1, aly-3, rsp-6, K08D12*.*3, R06C1*.*4, C53H9*.*2, rsp-5, ZK228*.*4, rpl-26*, and *pqn-70*), for which we could unambiguously identify the NMD eliciting isoform, and distinguish it from any other isoforms at the locus. All genes in this list were identified *de novo* (Fig 2E), as well as in multiple (≥ 2) previously published datasets.

### Vizualizations

All plot visualizations were produced with Python using publicly available libraries: plotly (v5.11.0), seaborn (v0.12.1), pandas (v1.5.2), and matplotlib (v3.6.2 - The axes join function needed by the coverage plotting scripts is depreciated and will be removed in v3.8+).

### Code availability

All scripts used are available on Github: https://github.com/MViscardi-UCSC/nanoporePipelineScripts.

**Figure S1: Distribution of poly(A) tail lengths for 17 endogenous NMD targets** Analysis performed as in Fig 3B on wildtype, *smg-5*, and *smg-6* animals between three replicates. Omission of a particular transcript species is due to insufficient read number.

**Figure S2: Cleavage of *ets-4* and *xbp-1* in *smg-6* and *smg-5* animals** Cleavage sites and respective nucleases for *smg-6* and *smg-5* libraries, diagrammed as in Fig 1B.

**Figure S3: Coverage plots of NMD targets and non-targets across 7 endogenous example genes** Analysis performed as in Figure 2C on wildtype, *smg-5*, and *smg-6* libraries across three replicates. From top to bottom, coverages are for the following categories: cleaved NMD isoforms (light green), full-length NMD isoforms (dark green), cleaved non-NMD isoforms (light blue), full-length non-NMD isoforms (dark blue), cleaved ambiguous isoforms (light gray), and full-length ambiguous isoforms (dark gray).

**Figure S4: Poly(A) tail length scatter plots for replicate libraries** Analysis performed as in Fig 3D, E and Fig 4C, D for replicate set 3.

**Figure S5: Fisher’s exact test analysis for replicate libraries** Analysis performed as in Fig 2E and Fig 5B, C for replicate set 3. Omission of a gene is due to insufficient read counts for that gene.

