## Supplementary figures and images for "NMD targets experience deadenylation during their maturation and endonucleolytic cleavage during their decay"

### Supplementary Figures

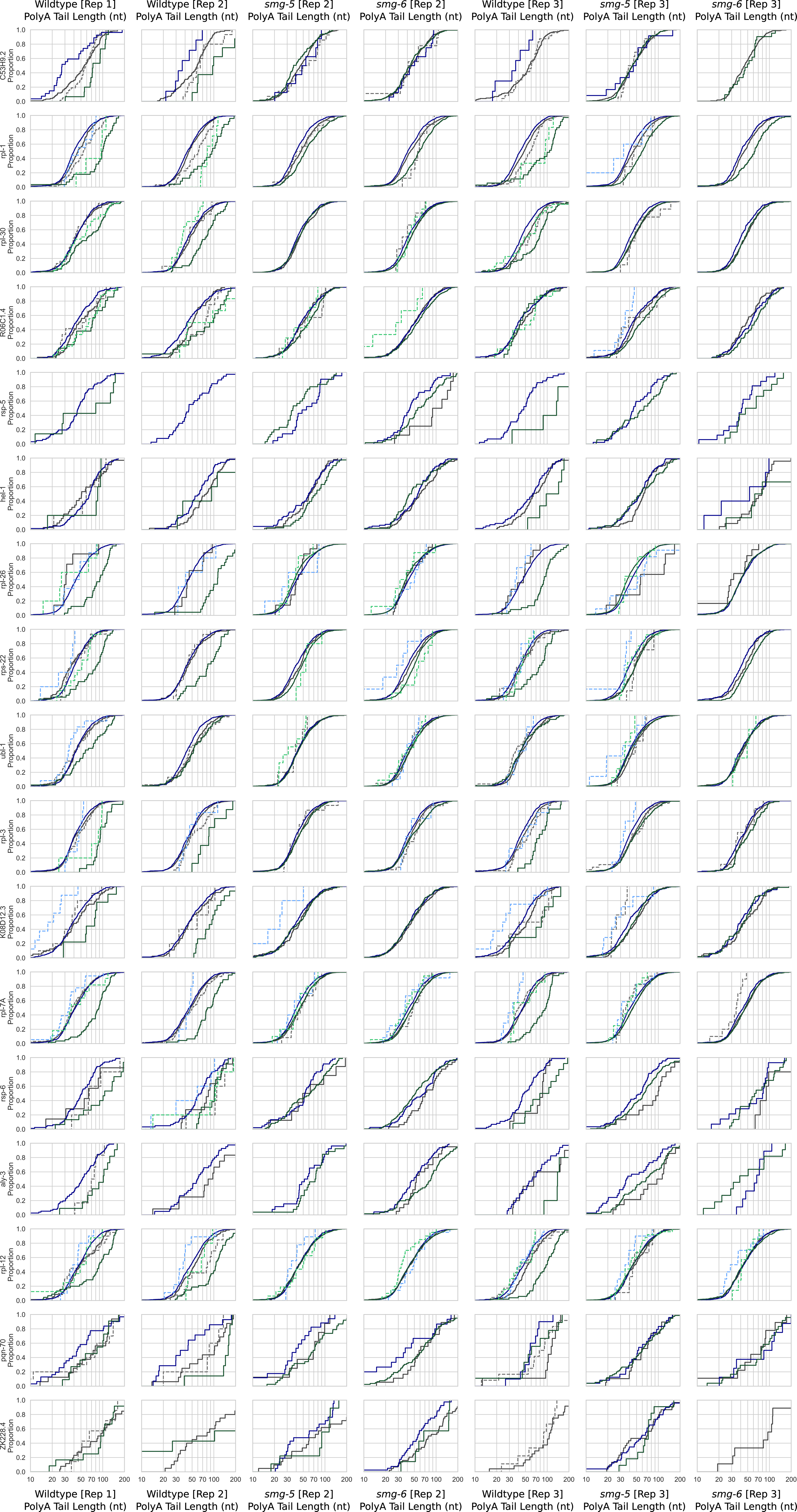

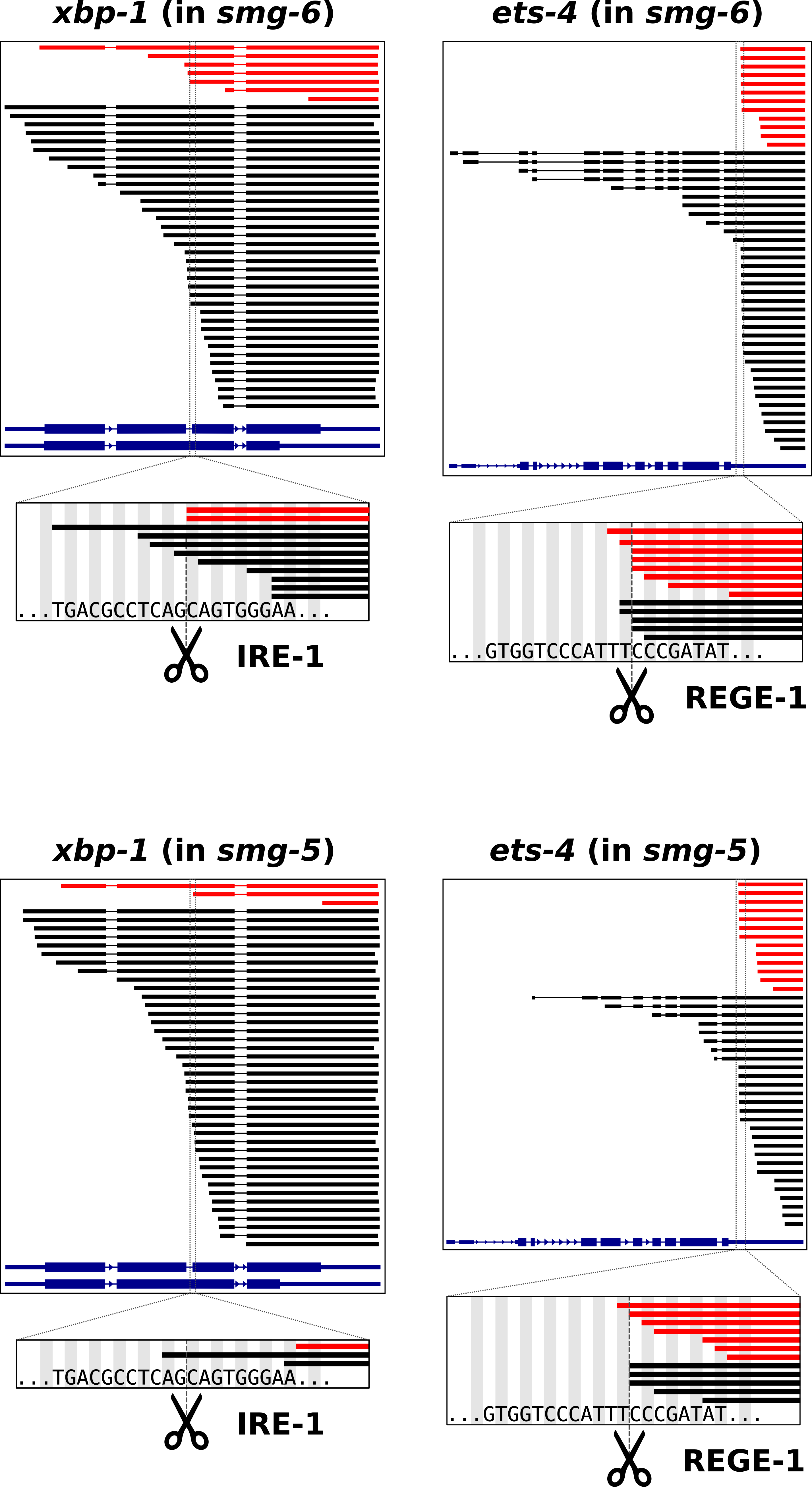

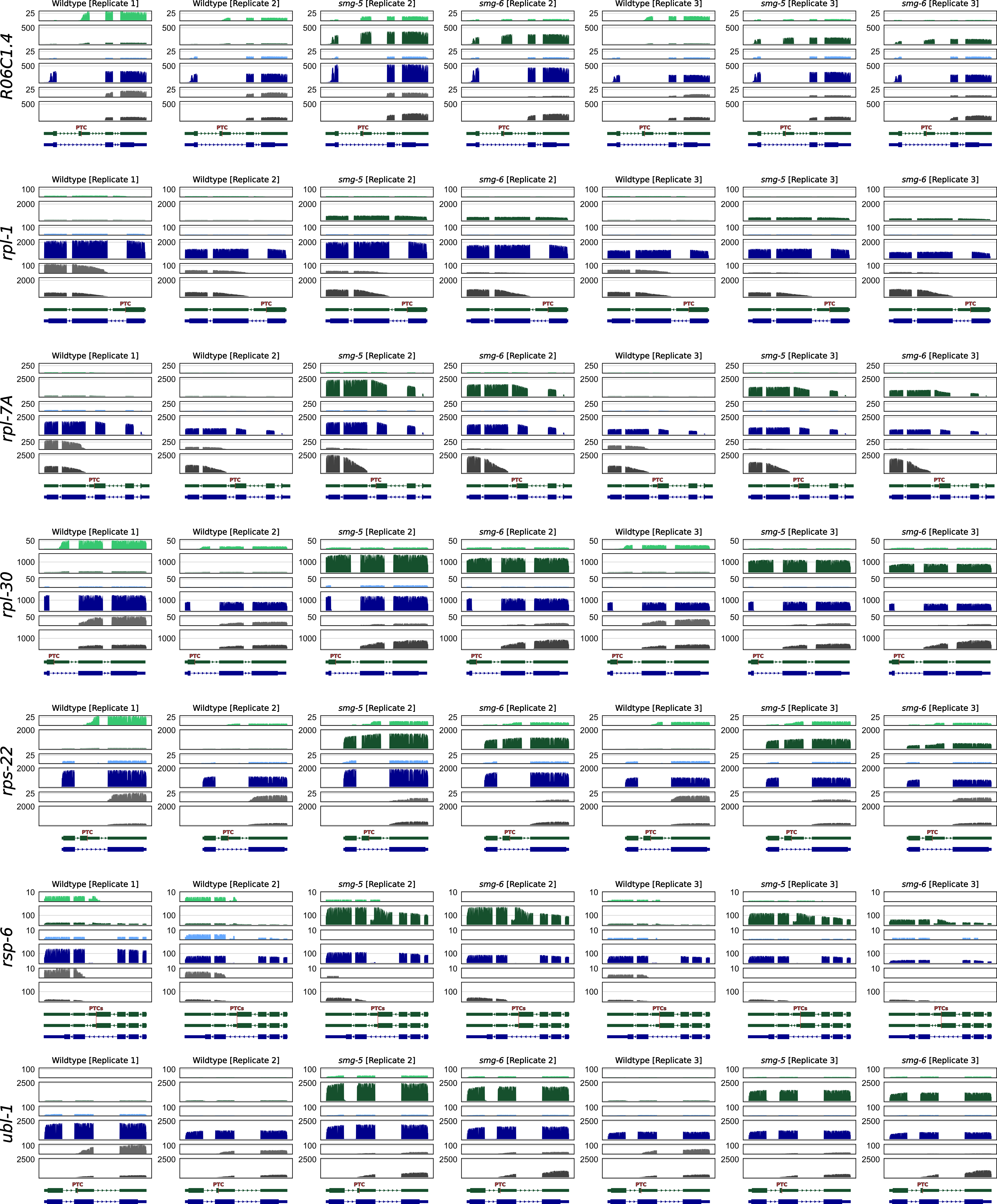

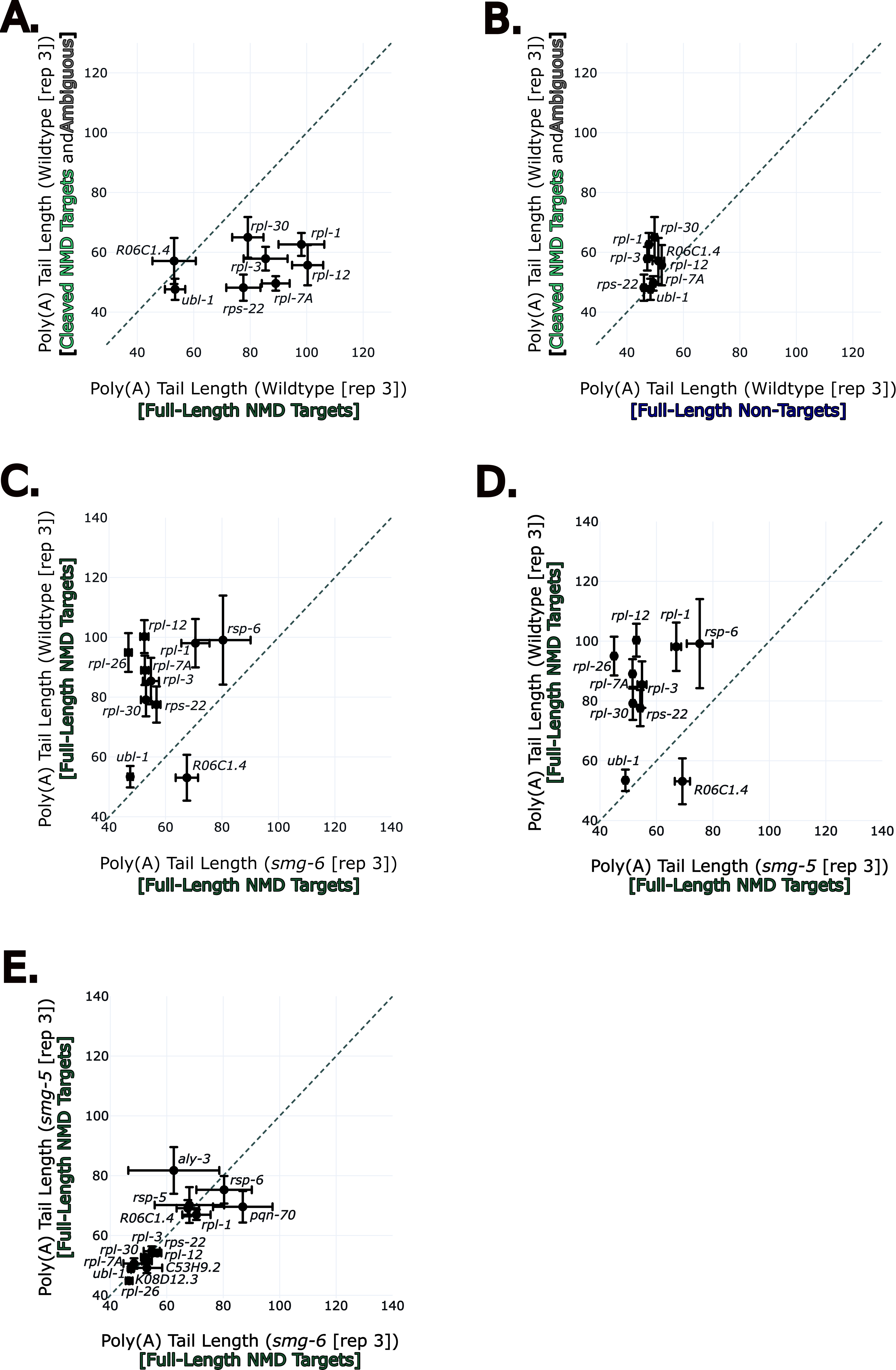

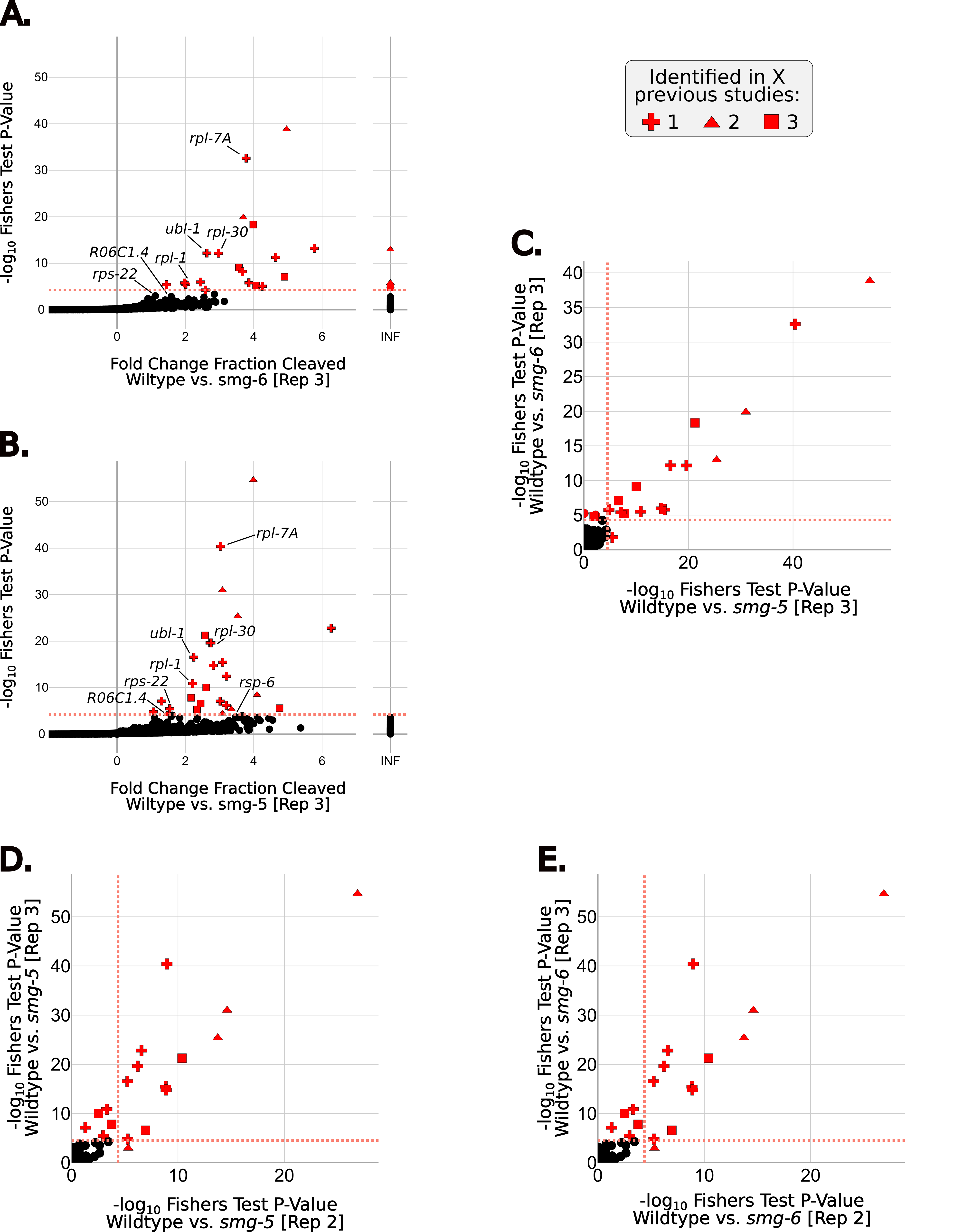
